# DNA methylation landscapes of matched primary and recurrent high grade serous ovarian cancers are preserved throughout disease progression and chemoresistance

**DOI:** 10.1101/2020.08.25.267161

**Authors:** Nicole Gull, Michelle R. Jones, Pei-Chen Peng, Simon G. Coetzee, Tiago C. Silva, Jasmine T. Plummer, Alberto Luiz P. Reyes, Brian D. Davis, Stephanie Chen, Kate Lawrenson, Jenny Lester, Christine Walsh, Bobbie J. Rimel, Andrew J. Li, Ilana Cass, Yonatan Berg, John-Paul B. Govindavari, Joanna K.L. Rutgers, Beth Y. Karlan, Benjamin P. Berman, Simon A. Gayther

**Author notes:** These authors contributed equally to the study (Nicole Gull, Michelle R. Jones, Pei-Chen Peng). These authors co-directed the study (Beth Y. Karlan, Benjamin P. Berman, Simon A. Gayther).

## Abstract

Little is known about the role of global DNA methylation in recurrence and chemoresistance of high grade serous ovarian cancer (HGSOC). We performed whole genome bisulfite sequencing (WGBS) and whole transcriptome sequencing (RNA-seq) to establish methylation and gene expression signatures in 62 primary and recurrent tumors from 28 patients diagnosed with stage III/IV HGSOC. Eleven of these patients carried pathogenic germline *BRCA1/BRCA2* mutations. Genome-wide methylation and transcriptomic features identified in primary tumors were largely preserved in matched recurrent tumors from the same patient (P-value = 7.16 × 10^−7^ and 1.41 × 10^−3^ in *BRCA1/2* and non-*BRCA1/2* cases respectively). Tumors from *BRCA1/2* carriers displayed high levels of heterogeneity, with significantly more shared methylation changes identified between primary and recurrent tumors from non-*BRCA1/2* patients, which may be related to the poorer survival we observe in HGSOCs from non-*BRCA1/2* carriers (P-value = 0.0056). Partially methylated domains (PMDs) dominated the epigenetic variation across all tumors, and were more hypomethylated in *BRCA1/2* than non-*BRCA1/2* cases. Differential gene expression analysis identified upregulation of genes from immune pathways including antigen processing and presentation in tumors from *BRCA1/2* carriers, implicating increased immune response in the improved survival observed in these patients. In summary, this study shows a previously unreported conservation of methylation and gene expression in recurrent HGSOCs. These data have implications for the possible effectiveness of epigenetic based therapies to treat both primary and recurrent ovarian cancers.

## INTRODUCTION

There is irrefutable evidence that methylation, as a mechanism of regulating gene expression, plays a key role in the development of most solid tumors. Germline or somatic hypermethylation is an alternative mechanism to pathogenic loss of function mutations caused by coding and splice site mutations, deletions and rearrangements that lead to allele specific gene deregulation. Hypermethylation silences genes that are critical components of genome integrity (e.g., DNA repair genes, cell cycle regulation genes and tumor suppressor genes) and may be an early event in the progression to cancer. Indeed, loss of expression of genes involved in cancer development occurs about 10 times more frequently by DNA hypermethylation of promoter CpG islands than through mutation of DNA^1–3^.

Early studies focused on the role of methylation within CpG islands of gene promoters as a mechanism to silence gene expression^4–6^. It has since become clear that methylation is a genome wide phenomenon that also targets the promoters of functional non-coding RNAs^7,8^, and more distal regulatory elements such as enhancers^9,10^. Thus, DNA methylation can also contribute to aberrant gene expression by altering the activity of transcription factor binding sites within enhancers and critical networks of gene expression variation involved in disease pathogenesis. More recent studies have shown that a global loss of methylation occurs in cancers, likely as part of the mitotic clock, across broad regions of the genome, known as Partially Methylated Domains (PMDs)^9,11–14^. These regions generally harbor genes expressed at low levels and account for the bulk of methylation changes that occur in cancer^9,11,12,14,15^.

Array based methods have become commonplace tools to evaluate the contribution of global methylation to tumor pathogenesis. As our knowledge of the landscapes of CpGs throughout the human genome has improved, so the content of methylation arrays have increased in scope and scale. However, even the latest arrays measure the methylation status of only a small fraction of the nearly 30 million known CpGs throughout the genome (e.g., the latest iteration, the Illumina MethylationEPIC BeadChip array covers 850,000 CpG sites). Recent advances in whole genome bisulfite sequencing (WGBS) for methylation profiling provide single-base resolution, expanding our ability to identify functionally relevant DNA methylation regions on the basis of transcriptional regulation. WGBS analyses have not yet been performed in substantially large numbers of tumors, but the method is already providing novel insights into the role of methylation in cancer.

The role of methylation in tumor recurrence and chemoresistance remains poorly understood. Despite early indications that demethylating agents may be effective treatments for high grade serous ovarian cancer (HGSOC)^16^ recent clinical trials of hypomethylating agents, including guadecitabine, have not yielded improvements in survival or resensitization to platinum-based therapies^17,18^. HGSOC is the most common and lethal histotype of ovarian cancer. About 70% of affected women are diagnosed with advanced stage disease (stages III/IV) and of these women <30% will survive more than five years. Patients are treated with maximal debulking surgery followed by combination chemotherapy with platinum. Typically, patients initially respond well to treatment, but usually relapse with recurrent and eventually chemoresistant disease^19^. Between 15% and 25% of tumors are classified with primary resistance^20^, and this tends to occur in patients with homologous recombination proficient tumors and/or amplification of the *CCNE1* locus at 19q12^21,22^. Nearly a third of all HGSOC cases have documented germline or somatic alterations in the *BRCA1* or *BRCA2* genes^22^, which results in DNA double strand break repair deficiency and an accumulation of DNA double strand break damage as tumors develop. The goals of this study were: (1) to establish the underlying role of global methylation in the recurrence and chemoresistance of HGSOC; and (2) to identify the role of global methylation in the development of primary HGSOC in women with and without germline defects in the *BRCA1* and *BRCA2* DNA double strand break repair genes.

## METHODS

### Cohort Description

Fresh-frozen primary and recurrent high-grade serous ovarian cancer specimens and DNA from 28 individuals diagnosed with high grade serous adenocarcinoma were included; 11 of whom had deleterious *BRCA1* and/or *BRCA2* germline mutations and 17 of whom do not harbor any known high or moderate risk mutations for HGSOC (*BRCA1, BRCA2, RAD51C, RAD51D, BRIP1* and *FANCM*) identified in clinical genetic testing. All patients were diagnosed with stage III or stage IV disease and underwent primary optimal surgical cytoreduction (to less than 1 cm residual disease) prior to administration of combination chemotherapy with platinum and taxane between the years of 1990 and 2014. For each patient, detailed clinical data were available including clinical genetic testing results, dates of original diagnosis and each subsequent recurrence, treatments administered throughout their disease course, operative and pathology reports, and other clinicopathologic variables including other cancer diagnoses and comorbidities.

### Specimen Acquisition and Preparation

Twenty-eight (28) consented patients with matching primary and recurrent tumors with DNA were identified in the Cedars-Sinai Medical Center Women’s Cancer Program Biorepository (IRB #0901). Fresh frozen tumors were embedded in optimal cutting temperature (OCT) compound, bisected and mounted and two slides were made for hematoxylin and eosin (H&E) staining. All slides were reviewed by a single pathologist to identify regions enriched for epithelial carcinoma (avoiding tumor stroma), which are then collected in a single punch of approximately 50mg collected on dry ice. Each punch was divided into three pieces, two of which were used for genomic DNA (gDNA) extraction using the Machery-Nagel Nucleospin DNA Kit, and the third for RNA extraction using the Machery-Nagel Nucleospin RNA Kit. DNA samples were assayed for quality using the QuBit (Thermo Fisher Sci, CA) to measure the content of double stranded DNA and by running 1ul gDNA on a 1.5% agarose gel at 100V for 1 hour to confirm no fragmentation of material has occurred during the extraction. RNA was extracted using an isopropyl-alcohol:chloroform approach following the standard operating protocol published by the Prostate Cancer Biorepository Network^23^. RNA was quantified on the Qubit in RNA mode to measure the amount of high quality dsRNA within the sample, and then on the Agilent Bioanalyzer, where an RNA Integrity Number (RIN) score is generated, reflecting the quality (by concentration and fragment size) of the samples. Germline DNA was extracted from whole blood drawn at the time of debulking surgery after diagnosis with HGSOC. DNA was extracted with the Qiagen DNEasy Blood & Tissue Kit (Qiagen, Germantown, MD, USA) and quantitated with the Quant-IT dsDNA Broad Range kit on a QuBit (Thermo Fisher, Waltham, MA, USA).

### Whole Genome Bisulfite Sequencing (WGBS)

Our workflow for WGBS required a minimum 300ng of high quality gDNA, which was sheared to approximately 175-200 bp using a Covaris sonicator, and bisulfite converted using the EZ DNA Methylation-Lightning Kit (Zymo). Libraries were constructed using the Accel-NGS Methyl-Seq DNA Library Kit (Swift Biosciences, MI), and amplified using no more than 6 cycles of PCR. Libraries were sequenced to at least 30x coverage (on average each base is sequenced thirty times) on the Illumina HiSeq4000 in 150bp paired end mode. This approach generated approximately 400 million read pairs per library, with a bisulfite conversion rate greater than 99%.

### WGBS Data Processing

WGBS reads were aligned to the human reference genome (build GRCh38) using BISCUIT^24^. Duplicated reads were marked using Picard Tools^25^. Methylation rates were called using BISCUIT. CpGs with fewer than 5 reads of coverage were excluded from further analysis. Adapter sequences were trimmed using TrimGalore, using default parameters for Illumina sequencing platforms^26^. Quality control was performed using PicardTools as well as MultiQC^27^. Bisulfite non-conversion was checked using the Biscuit QC module in MultiQC^28^. Principal Component (PC) analysis was performed on CpGs with coverage ≥10 and the top 10,000 most variable CpGs were included in the identification of the top 10 PCs using the prcomp function from the stats package in R^29^.

### Calling Partially Methylated Domains (PMDs)

To call Partially Methylated Domains (PMDs), we first divided the genome of each sample into non-overlapping 100kb bins, and took the average of all solo-WCGW CpGs within each bin, using the solo-WCGW definition from Zhou et al.^30^. We then converted the methylation averages to M-values (Mi=log2(Betai/(1-Betai))^31^, and fit M-values to a 3-component Gaussian Mixture Model (GMM) using the mclust R package^32^. Based on its mean, we assigned the three components labels of low, intermediate, and high. Each bin was labeled as a PMD if the probability of being in the high bin was less than 0.01, and multiple consecutive PMD bins were merged into a single PMD call. Common ovarian cancer PMDs (ovcaPMDs) were defined as regions identified from PMDetect that overlapped between 9-19 of our samples. Common ovca PMDs were combined with common PMDs from other cancer cell types^30^. Solo-WCGW scores were calculated by averaging the methylation of solo-WGCW CpGs (using the definition from Zhou et al.^30^) within the combined ovcaPMD+commonPMD set. Taken together these PMDs spanned 69.57% of the genome, comprising 14.96% of the genome spanned by ovcaPMDs and 54.6% of the genome spanned by the common PMD set.

### Calling Differentially Methylated Regions (DMRs)

PMDs were masked from each sample’s BED file before conducting differentially methylated region (DMR) analysis. The Bioconductor package dmrseq^33^ was utilized to identify DMRs between *BRCA1/2* carriers and *BRCA1/2* non-carriers using default settings. Metilene^34^ was used to identify DMRs between matched primary and recurrent tumors from each patient using the following parameters: -M 500, -m 5, p 0.1, -c 5. Only DMRs that overlapped between two or more patients were retained. Bedtools^35^ merge function was performed on all overlapping DMRs to merge regions within 250bp using parameter -d 250. Heatmaps were plotted using mean methylation across each identified DMR. Enrichment analysis of DMRs was conducted using annotatr^36^ and ChIPseeker^37^. Backgrounds for enrichment controlled for either DMR size only or DMR size plus CpG count. A DMR size only background was generated using bedtools shuffle^35^. DMR size plus CpG count background was generated using an in-house developed script.

### RNA-Seq Library Preparation and Sequencing

RNA was extracted using the protocol published online by the Prostate Cancer Biorepository network (SOP#006), where frozen tissue was stored at −80C until extraction. 1mL of trizol was added to the tissue in a 1.5mL eppendorf tube and incubated for 5 mins at 15-30C to dissociate nucleoprotein complexes. Next 200ul of chloroform was added and tubes capped and mixed vigorously for 15 secs then incubated at room temperature for 3 mins. Samples were then centrifuged for 15 mins at 4C at 12,000g. The top, aqueous phase was removed to a fresh, sterile 1.5ml tube and mixed with 500ul of isopropyl alcohol to precipitate the RNA. Samples were then incubated at room temperature for 10 mins and then centrifuged for 10min at 4C at 12,000g. A pellet was visible in high yield samples, and supernatant was removed (and discarded), leaving the pellet untouched. The pellet was washed with 1mL 75% ethanol and mixed by vortexing and then centrifuging at 7,500g for 5min at 4C. The supernatant was removed (leaving the pellet untouched) and pellet allowed to air dry before being dissolved in 200ul RNase-free water and incubated at 55C for 10 mins. Sample concentration was measured using the Qubit RNA Broad Range kit and sample quality was measured using the Agilent BioAnalyzer 2100. Sequencing libraries were prepared by adding 1ug of RNA to the TruSeq Stranded Total RNA Kit with Ribo- and Mito-depletion following the TrSeq standard protocol with 15 cycles of PCR. Libraries were quantified using Qubit DNA Broad Range kit and pooled before being run on one lane of a HiSeq2000 to collect ~1M reads per library for quality control. PCR duplication rate was estimated in this low coverage sequencing run in 150bp paired end mode, and library complexity was estimated using PreSeq. Based on the complexity measured in this low coverage sequencing experiment we estimated the maximal coverage that would continue to provide informative measurement of transcripts in the library was ~350M reads. Each library was then pooled and this pool was sequenced in 2×150bp mode on an Illumina Novaseq 6000, and we generated ~335 million reads from each library. Our data analysis workflow for RNA-Seq has been developed specifically to improve gene feature identification and measurement in archived frozen tissue samples, which can perform poorly using standardized workflows.

### RNA-Seq Quantification and Statistical Analysis

Reads within each fastq file were first trimmed using TrimGalore to remove low quality bases and sample barcodes, retaining reads 75bp or longer. Each transcriptome is then aligned to hg38 and the Gencodev29 primary assembly^38^. Genes are quantified with RSEM^39^ and Kallisto^40^. Sample-specific gene models were generated using alignments produced with STAR two pass mapping and Stringtie^41^. Gene expression values were shown as normalized variance stabilizing transformation (vst) counts. To measure RNA abundance, we first obtained BAM alignment quality metrics using Picard (http://broadinstitute.github.io/picard). Samples with less than 90 percent of reads mapped to the correct strand of the reference genome (PCT_CORRECT_STRAND_READS) were omitted. Patients whose primary and recurrent tumors both passed this quality control were retained (n=50) (**Supplementary Table 2**). Read counts were quantified using the R package Salmon^42^ at transcript level and reads were mapped to Genecode Release 29 (GRCh38) comprehensive gene annotations by R package ‘tximport’^43^. To filter out potential artifacts and very low expressed transcripts we retained transcripts with length greater than 300 bp, TPM (Transcripts Per Million) value greater than 0.05 and isoform percentage greater than 1%. Transcripts in blacklist regions^44^ were also filtered out. We retained transcripts expressed in more than 5 samples, which resulted in 91,411 transcripts from 33,969 genes. Tumor purity was estimated by the degree of heterogeneity of the tumor microenvironment. We applied the R package ‘consensusTME’^45^ to estimate cell type specific enrichment scores based on TCGA ovarian cancer data, and confirmed there was low stromal content and generally low infiltration by immune cells across the samples (**Supplementary Figure 6**).

Differentially expressed genes between the *BRCA1/2* carrier versus *BRCA1/2* non-carrier and primary versus recurrent tumors were detected by R package ‘DESeq2’^46^. The P-values were adjusted for multiple testing using the Benjamini-Hochberg procedure. Since our experiment design has group-specific effects, comparisons between *BRCA* carrier status are made between patients, while comparisons between primary versus recurrent tumors are made within the patient. To control for confounding differences between the primary and recurrent tumors from patients we constructed a nested DEseq2 model with formula; ~purity + BRCAStatus + BRCAStatus:PatientID + BRCAStatus:isRecur, which has the main effect for BRCA status plus nested interactions with primary and recurrent status. To see whether the identified gene sets (e.g. genes inside PMDs or differentially expressed genes) show significant functional concordance, we performed Gene Set Enrichment Analysis ^47^ for Kyoto Encyclopedia of Genes and Genomes (KEGG) pathways^48–50^. We implemented enrichment analysis with R package ‘clusterProfiler’ ^51^. For each enrichment analysis, we set the number of permutations to 10,000 and reported enriched pathways with a Benjamini-Hochberg adjusted P-value less than 0.05.

### Linking enhancers to target genes

Correlation between DMRs and gene expression was performed by comparing primary and recurrent tumors from each individual patient, for samples with matched WGBS and RNA-seq available for both primary and recurrent tumors (n = 28). Only DMRs that overlapped between two or more patients were included in this analysis. Using GENCODE28, DMRs regions >2kb from any TSS were annotated as “distal” and regions <2kb from TSS were annotated as “promoter”. Distal regions were mapped to the closest genes (10 upstream and 10 downstream) and the promoter to the closest gene and the correlation between their expression and methylation measured, where average beta value of the DMR correlates (using Spearman test) to a change (positive or negative) in expression of the nearby genes. ELMER version 2.8.3^52^ was used to map the genes, and the correlation was performed to each link (DMR-gene) only using the samples in which the DMR was identified using the function cor.test in R. Links with a minimum P-value of 0.05 were retained.

## RESULTS

### Whole genome bisulfite sequencing (WGBS) in primary-recurrent high grade serous ovarian cancers

We used WGBS to perform whole genome methylation profiling and RNA sequencing (RNA-seq) for whole transcriptome profiling of 62 fresh frozen primary or recurrent tumor tissues from 28 women diagnosed with stage III/IV high grade serous ovarian cancer (HGSOC). All patients were treated at Cedars Sinai Medical Center. Clinical features of the patients and their tumors are given in **Supplementary Table 1** and illustrated in **Figure 1**. Eleven of the 28 patients carried a germline, pathogenic mutation in the *BRCA1* and/or *BRCA2* genes; the remaining 17 patients were confirmed non-*BRCA1/2* germline mutation carriers (**Supplementary Table 1**). All patients received similar first line treatments comprising optimal debulking surgery followed by combination chemotherapy with a platinum agent. Time to first recurrence and median survival times were significantly greater in *BRCA1/2* carriers compared to non-*BRCA1/2* carriers (2768 days vs 1678 days respectively, P-value=0.0056) which is consistent with previous studies (**Figure 1b, 1c**)^22,53,54^.

**Figure 1.**
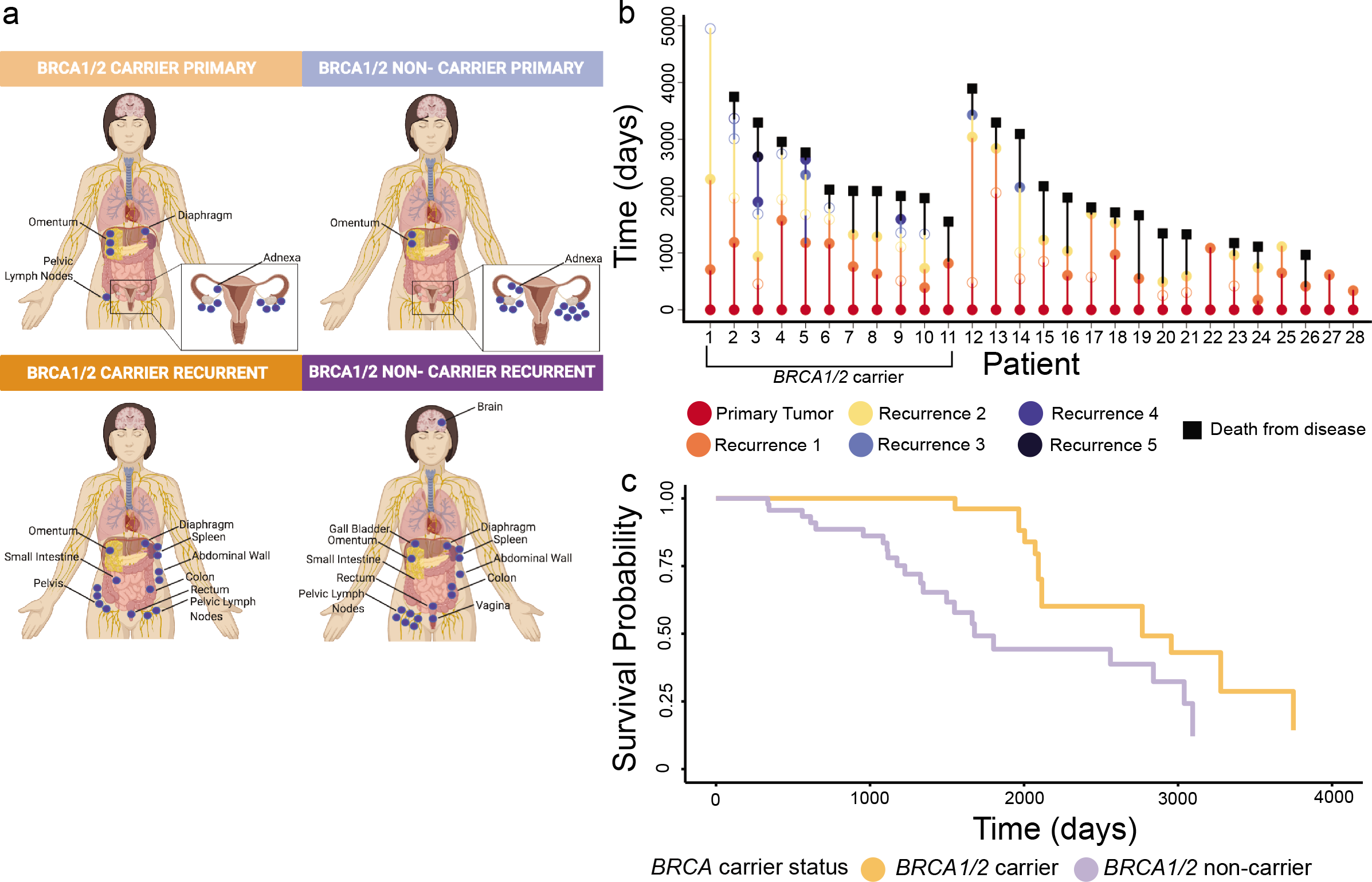
Clinical features of HGSOC patients with recurrent disease. **(A)** Tumor location of primary and recurrent tumors in women with and without germline *BRCA1/2* mutations. **(B)** Disease course for each patient profiled in the cohort. **(C)** Women with *BRCA1/2* mutations have an improved survival, even with recurrent disease

For each specimen, we generated a bisulfite converted library and sequenced to a depth of ≥30x, generating approximately 400 million read pairs per library, with a bisulfite conversion rate greater than 99%. After removing CpGs covered by fewer than 5 reads, we obtained on average of 24.2 million CpGs covered per tissue specimen (range 13.1 - 26.5 million), with an average of 24.6 million CpGs in primary tumors (15.9 - 26.5 million) and 23.8 million CpGs in recurrent tumors (13.1 - 26.4 million). In addition, RNA was extracted from frozen tissue samples and sequenced, which generated about 335 million reads from each library. After removing very low expressed transcripts and transcripts in blacklist regions, we obtained 91,411 transcripts from 33,969 genes.

### The methylation landscape of primary and recurrent HGSOCs is preserved in individual patients but highly heterogeneous between patients

By evaluating genome wide maps of all CpG locations across each chromosome (**Figure 2a**) and through additional analysis of the 10,000 most variable CpGs across all tumors (**Supplementary Figure 1a**), we observed widespread variability in methylation profiles between different patients. Some regions showed evidence of variable methylation between primary and recurrent tumors (e.g., hypomethylation at exon 2 of *GJC2* in tumors, **Figure 2b**), but generally the methylation profiles of primary tumors were maintained in recurrences from the same patient (**Supplementary Figure 1a**).

**Figure 2.**
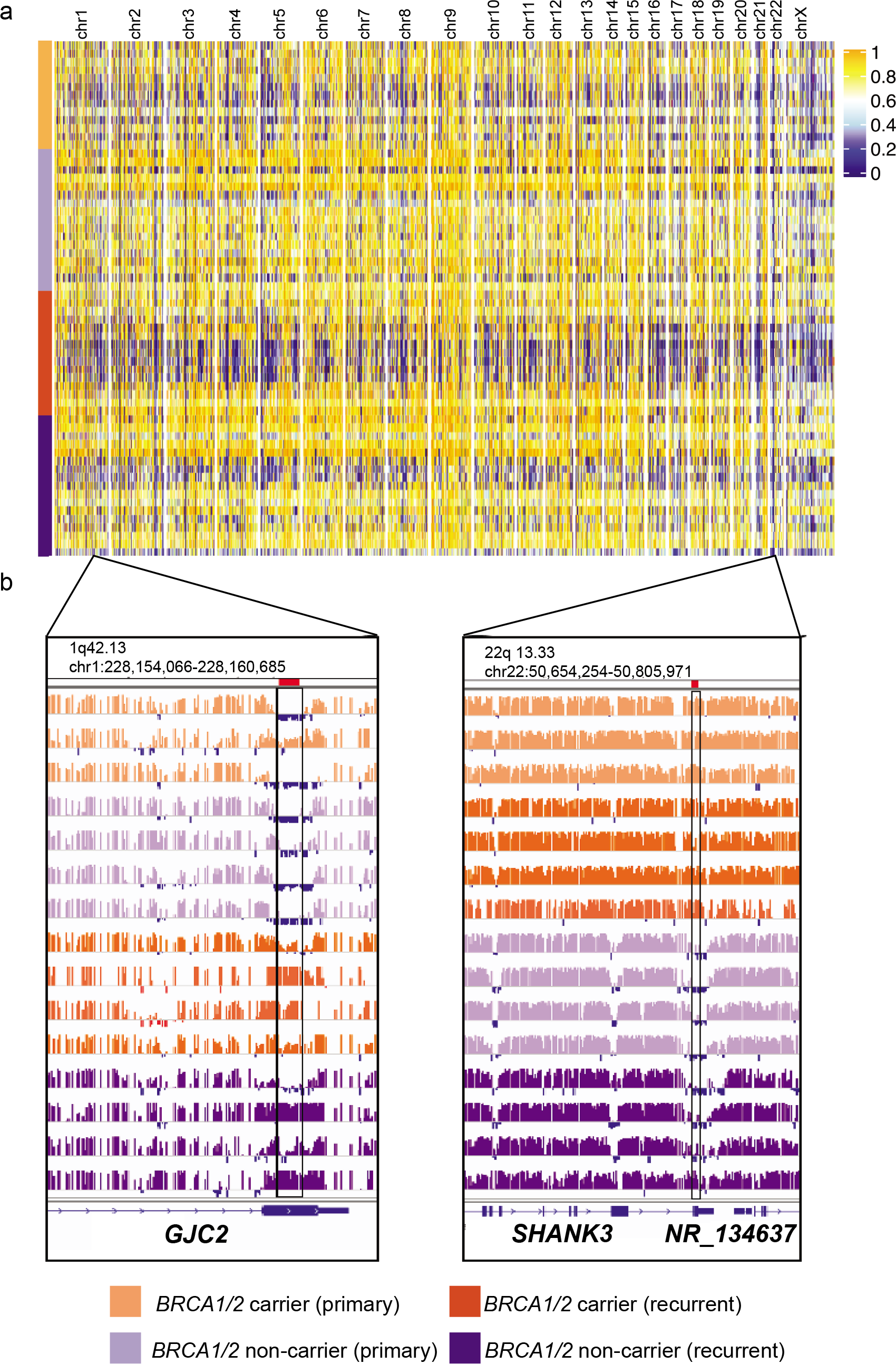
High grade serous ovarian cancers show heterogeneous patterns of genome-wide methylation. **(A)** Primary and recurrent tumors show heterogeneous patterns of methylation across the genome, with many tumors showing extensive hypomethylation on the X chromosome. CpG values are averaged across 10kB windows, minus ENCODE blacklist regions **(B)** Examples of two regions on chromosome 1q42.13 and 22q13.33 showing differentially methylated regions (boxed regions) from two comparisons - Primary vs Recurrent tumors (left) and *BRCA1/2* carrier vs *BRCA1/2* non-carrier (right)

Cancers often undergo a global loss of methylation within large genomic blocks (partially methylated domains or PMDs), and so we characterized the landscapes of PMDs in each primary and recurrent tumor. The fraction of the genome covered by PMDs varied between tumors, from less than 1% up to 58%. On average 29% of the genome was covered by PMDs in our sample set (**Supplementary Table 3**). There was no consistent pattern in the PMD architecture across this series of tumors; less than 0.03% of PMDs were shared in 56/62 tumors (**Figure 3a**). We identified a set of ‘common’ PMDs that were shared in at least 9 tumor specimens (ovcaPMDs), determined by the first inflection point of the bimodal distribution seen by plotting PMD frequency (**Supplementary Table 4**).

**Figure 3.**
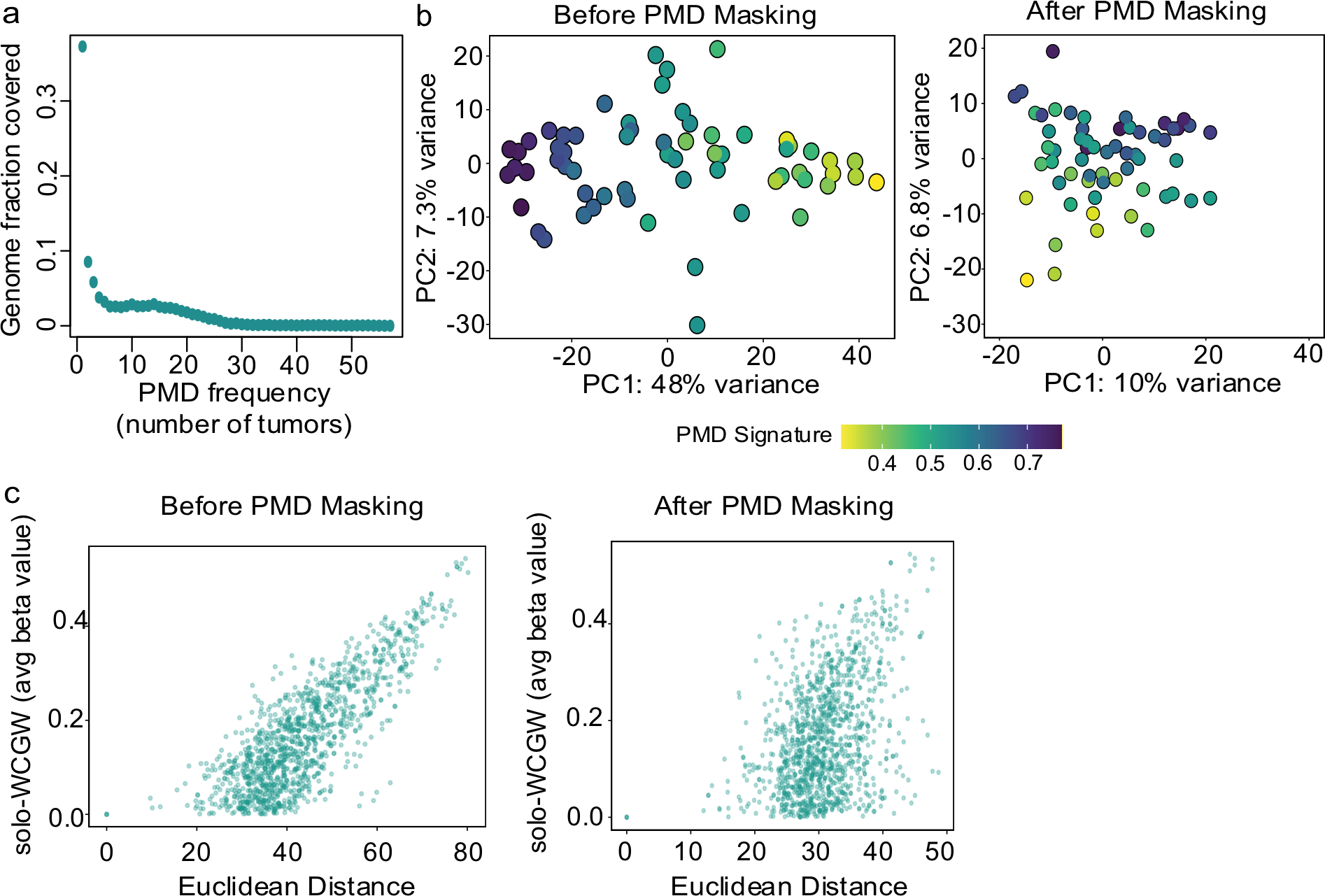
Hypermethylation within PMDs in high grade serous ovarian cancers is driven by soloWCGWs. **(A)** Most PMDs detected across the cohort were unique to a single tumor, with only 2% of PMDs observed in more than 30 tumors. **(B)** Principal components (PC) analysis identifies a large proportion of the variance between tumors was due to methylation at soloWCGW sites within PMDs. Masking the genome for common PMDs and ovcaPMDs removed much of the variance. After masking tumors clustered by patient, rather than by germline mutation status or disease stage. **(C)** The strong correlation between PMD soloWCGW values and pairwise Euclidean distance between tumors was lost after masking PMDs

In order to investigate the global differences between tumors and the influence of PMD associated hypomethylation, we performed Principal Component (PC) analysis for each tumor using the top 10,000 most variable CpGs (**Figure 3b**). The first PC dominated the clustering, accounting for 48% of total variance. PMD hypomethylation was shown to have both strong sequence as well as regional effects, with “soloWCGWs” (i.e. CpGs flanked by A/T on either side and no other CpGs within a 150bp window) having the strongest hypomethylation^30^. When we calculated average soloWCGW methylation within a maximal set of PMDs that included both cell-type ‘invariant’ PMDs described by Zhou and colleagues^30^ and the set of common regional ovcaPMDs, PC1 was almost completely correlated with soloWCGW PMD methylation (**Figure 3b, left**). When we removed from the analysis CpGs that were covered by the combined common PMD/ovcaPMD set and re-performed PCA analysis, the variance included in PC1 was reduced from 48% to 10% and the overall association with soloWCGW PMD methylation was weaker (**Figure 3b, right**), indicating that methylation in PMD regions contributes significantly to the overall appearance of heterogeneity across this set of tumors. In order to remove the influence of highly variable PMD methylation, we hereafter used the PMD-masked regions of the genome in performing the analysis of differentially methylated regions (DMRs). We also checked the effects of masking by performing an all vs. all pairwise comparison of soloWCGW methylation difference vs. overall Euclidean distance (calculated based on the 10,000 most variable CpGs), for all tumor samples (**Figure 3c**). As expected, the correlation was significantly removed by PMD masking (r^2^=0.65 before masking and r^2^=0.17 after masking).

In order to investigate the global methylation patterns that remained, we used the 10,000 most variable post-masking CpGs to perform hierarchical clustering analysis. Even after removing the global effects of PMD hypomethylation, primary and recurrent tumors from the same patient clustered together, independent of *BRCA1/2* mutation status, tumor site or experimental batch (**Supplementary Figure 1a**). We quantified this by comparing Euclidean distances from all tumor pairs from the same patient (intra-patient distances) vs. the distances of all pairs from different patients (inter-patient distances). Intra-patient distances were universally smaller than inter-patient differences, with average intrapatient distances of 21.78 and 23.45 respectively in *BRCA1/2* and non-*BRCA1/2* carriers, and average inter-patient distance of 31.69 and 30.76 respectively in *BRCA1/2* and non-*BRCA1/2*: P-values = 7.16 × 10^−7^ and 1.41 × 10^−3^ (**Figure 4a**). We performed the same analysis for global RNA-seq profiles, with almost the same results (average intra-patient distances of 153.15 and 149.72, and average inter-patient distance of 192.93 and 194.03: P-values = 9.67 × 10^−5^ and 6.70 × 10^−4^ in *BRCA1/2* carriers and non-*BRCA1/2* carriers respectively; **Figure 4a**). The results from DNA methylation and RNA-seq analyses are in agreement that the functional epigenetic landscape of recurrent tumors is dominated by inter-patient heterogeneity rather than disease progression, and the validation by RNA-seq demonstrates that the DNA methylation differences are not the result of residual PMD hypomethylation differences that remain after masking.

**Figure 4.**
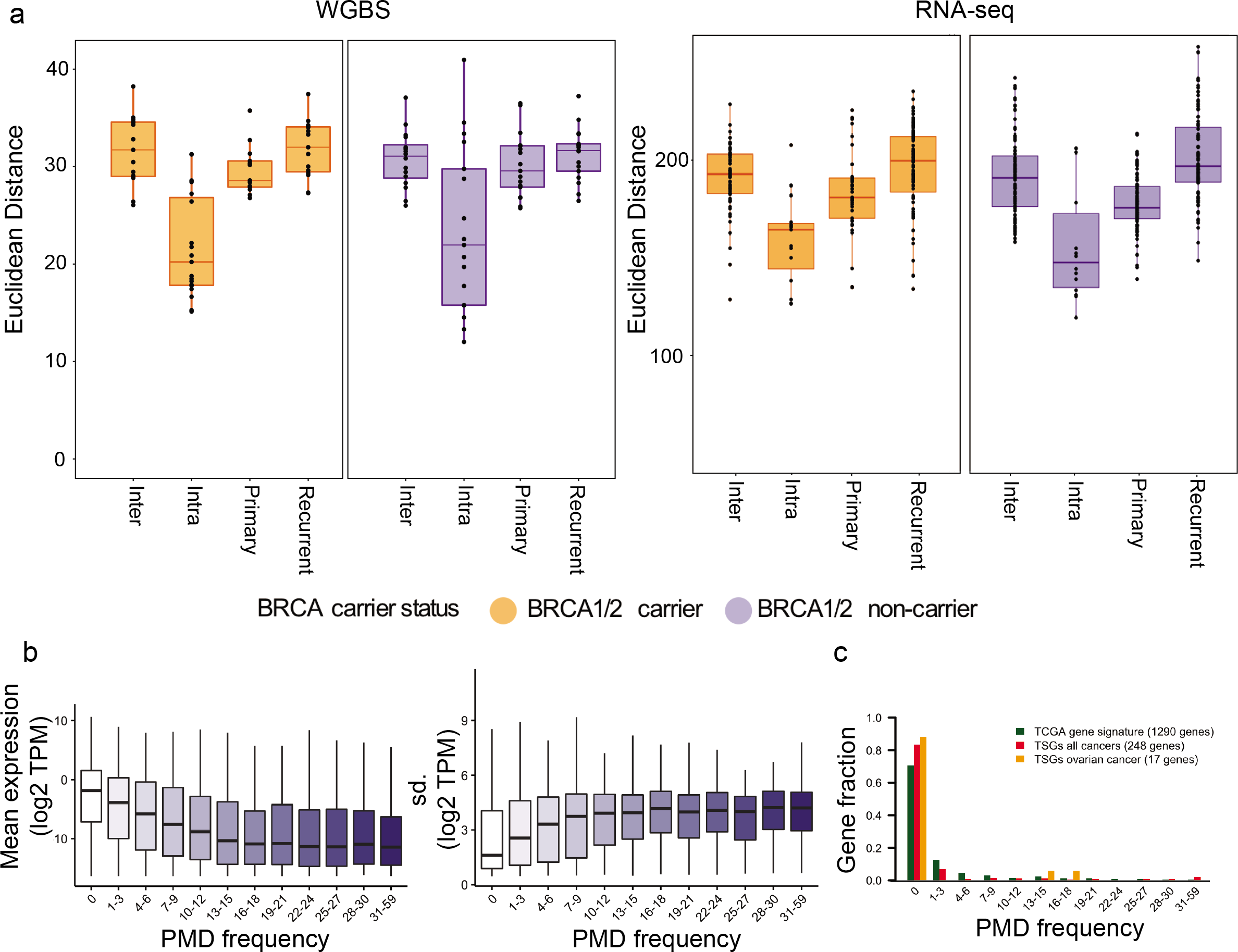
Methylation and transcription are largely preserved between primary and recurrent tumors from each patient. Expression of genes within PMDs. **(A)** Intrapatient pairwise Euclidean distances were significantly smaller than inter-patient distance or the intra-stage stage distance in both methylation (top) and gene expression (below) from paired RNA-Seq. **(B)** Genes within PMDs are less expressed (left) but more variable in their expression (right) than genes outside of PMDs. **(C)** The vast majority of tumor suppressor genes TSGs in cancer and that form the ovarian cancer molecular subtypes defined by TCGA are located outside of PMDs.

We observed that genes within PMDs were expressed at a lower level than genes outside PMDs, with lowest expression for genes inside highly recurrent PMDs (P-value <2 × 10^−16^, linear regression; **Figure 4b**, left panel). The variability in transcript expression for genes within PMDs was greater than for genes outside PMDs (P-value <2 × 10^−16^, linear regression; **Figure 4b**, right panel). These findings were similar in tumors with different *BRCA1/2* and *non-BRCA1/2* mutation status, and between primary and recurrent tumors (**Supplementary Figure 2**). Given PMDs mark repressive expression, we examined the frequency of known or predicted tumor suppressor genes (TSGs)^55^ located in our PMD set (common PMDs plus ovcaPMDs). Most TSGs (207/248, 83%) are located outside of PMDs with only 17/248 TSGs (7%) lying in low-frequency PMDs (hypergeometric test P-value = 4.17 × 10^−17^; **Figure 4c**). TSGs associated with ovarian cancer development also tended to lie outside PMDs (P-value=0.002), indicating they are likely to be more highly expressed. We also look at gene location with respect to PMDs for differentially expressed genes identified by the cancer genome atlas project (TCGA) associated with the different molecular subtypes of HGSOC^22^. Seventy-one percent of genes that comprise the subtype specific gene expression signatures of HGSOCs were located outside PMDs (P-value = 2.75 × 10^−18^). Gene Set Enrichment Analysis (GSEA) for genes located within PMDs shows that these genes are enriched in several signaling pathways, such as Rig-I-like receptor signaling and ErbB signaling pathways (**Supplementary Table 5**), which are similar to findings in breast cancer^14^ and human neuron cells^56^.

### Methylation changes are not consistently observed across recurrent high grade serous ovarian tumors

We evaluated differential methylation changes in non-PMD regions between primary and recurrent tumors to identify regions and/or specific molecular markers that may be associated with the development of recurrence after a primary diagnosis of HGSOC. We first performed differentially methylated region (DMR) analysis to identify methylation differences between the set of all primary tumors and the set of all recurrent tumors (using PMD-masked genome). There were 15,082 DMRs (Q-value < 0.1) across all primary and recurrent tumors but no statistically significant DMRs. There were significantly more DMRs in primary versus recurrent tumors from non-*BRCA1/2* carriers (11,205 significant DMRs in total, average of 659 DMRs per case comparison) compared to tumors from *BRCA1/2* carriers (3,877 significant DMRs in total, average of 388 DMRs per case comparison) (P-value = 0.004). We retained DMRs between primary and recurrent tumors that were shared between at least two patients and merged overlapping regions that were within 250bp of each other, yielding a total set 1,785 recurrent DMR regions (**Supplementary Table 6**). We plotted these regions as the delta, or overall change, in methylation levels at each CpG site identified within a DMR per patient group. Of these DMRs, 558/1785 (31%) were variably methylated (observed as hyper- and hypo-methylated in individual patients; **Supplementary Fig 3a**). Hierarchical clustering analysis of the 1,785 DMRs did not identify any clinical or molecular features that correlate with consistent methylation changes in two or more patients across the cohort (**Supplementary Figure 3a**).

To determine if there were any trends in PMD methylation in the progression from primary to recurrent cancer, we evaluated the average PMD soloWCGW methylation values in each tumor, but again found no common or consistent changes (**Supplementary Figure 3b**). Neither did we find enrichments of DMRs for specific functional genomic features or the expression of local genes. Ninety-nine genes were differentially expressed between all primary and recurrent tumors (adjusted P-value <0.05), of which 37 genes had lower expression and 62 genes higher expression in recurrent versus primary tumors (**Supplementary Table 7**). Differential expression for twenty of these genes was directionally consistent with differential changes in methylation (**Supplementary Table 8**). Taken together, these data indicate that the methylation landscapes of recurrent HGSOCs remain relatively stable compared to the primary tumors from the same patients, and do not acquire common somatic methylation changes that may be drivers of tumor recurrence/ chemoresistance.

### Differential methylation changes correlate with gene expression in *BRCA1/2* versus non-*BRCA1/2* tumors

We compared the patterns of methylation in primary and recurrent tumors from *BRCA1/2* and non-*BRCA1/2* mutation carriers. DMR analysis using DMRseq^33^ identified 135 significant DMRs in tumors identified between these groups (Q-value < 0.05; **Figure 5a**, **Supplementary Table 9**). PCA analysis identified a trend of differential methylation in tumors from *BRCA1/2* versus non-*BRCA1/2* carriers (**Figure 5b**). Tumors from *BRCA1/2* carriers were more hypomethylated compared with tumors from *non-BRCA1/2* carriers both in DMRs (101 DMRs hypomethylated in *BRCA1/2* carriers vs. 34 DMRs hypomethylated in non-*BRCA1/2* carriers) and in PMD regions (P-value = 0.0011) (**Supplementary Figure 4a**). As with previous analyses, DMRs between primary and recurrent tumors from the same patient were more conserved than between tumors from different patients (**Supplementary Figure 4b**). Annotation of DMRs hypermethylated in non-*BRCA1/2* tumors showed that these regions were enriched in GENCODE promoters and enhancers identified in ovarian cancer cell lines (**Supplementary Figure 5b**), indicating that specific genes in these tumors may be inactivated by hypermethylation through their associated regulatory elements. Regions hypermethylated in *BRCA1/2* tumors showed no promoter or enhancer enrichment (**Supplementary Figure 5a**).

**Figure 5.**
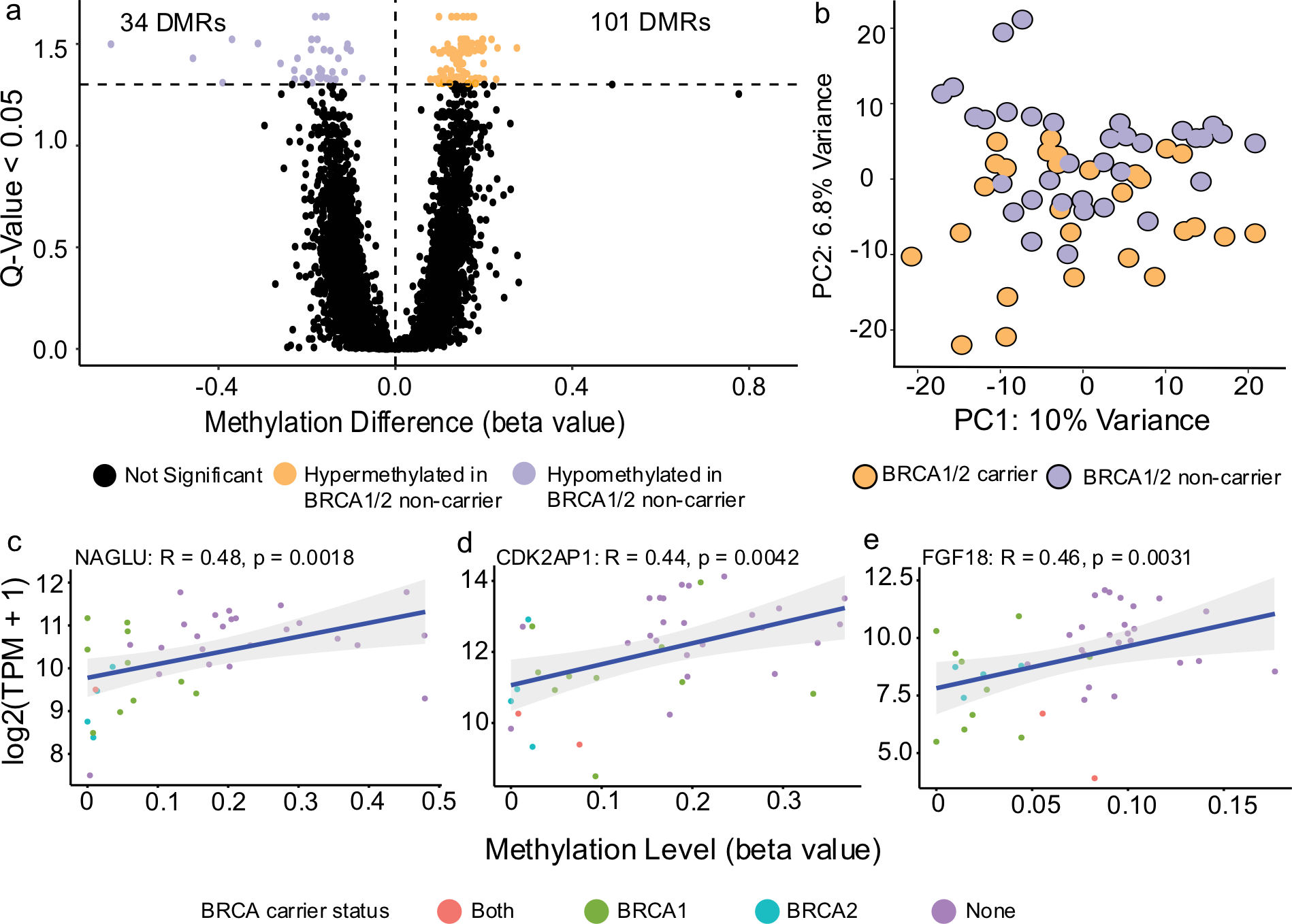
Differences in methylation by BRCA1/2 carrier status. **(A)** 135 differentially methylated regions (DMRs) were identified in tumors from *BRCA1/2* carriers compared to non-*BRCA1/2* carriers with a trend towards hypermethylation in *BRCA1/2* non-carrier tumors; 101 regions hypermethylated in *BRCA1/2* non-carriers compared to only 34 regions hypermethylated in *BRCA1/2* carriers **(B)** Principal components analysis of HGSOC tumors using PMD masked data shows a trend towards differences in the tumors based on carrier status. **(C-E)** Individual genes where methylation levels within hypermethylated regions were directionally correlated with gene expression; in order – NAGLU, CDK2AP1, FGF18

PCA analysis based on RNA-seq data from these tumors were consistent with methylation data (**Figure 6a**). There were 3,341 differentially expressed genes (DEGs) between *BRCA1/2* and non-*BRCA1/2* tumors (adjusted P-value <0.05) (**Supplementary Table 10**) of which 1,760 genes were up-regulated and 1,581 genes were down-regulated in *BRCA1/2* tumors (**Figure 6b**). The observation of more up-regulated genes is consistent with the directionality of increased activity/hypomethylation in *BRCA1/2* carriers. Up-regulated genes in *BRCA1/2* tumors were significantly enriched in immune related pathways including autoimmune diseases, infection response and antigen processing and presentation (**Figure 6c**) even though there were no noticeable differences in immune cell infiltration in these tumors compared to non-*BRCA1/2* tumors (P-value = 0.97). Down-regulated genes in *BRCA1/2* tumors were most significantly enriched in pathways that maintain stemness and cell differentiation, including the hippo signaling pathway (adjusted P-value <0.05) (**Figure 6c**).

**Figure 6.**
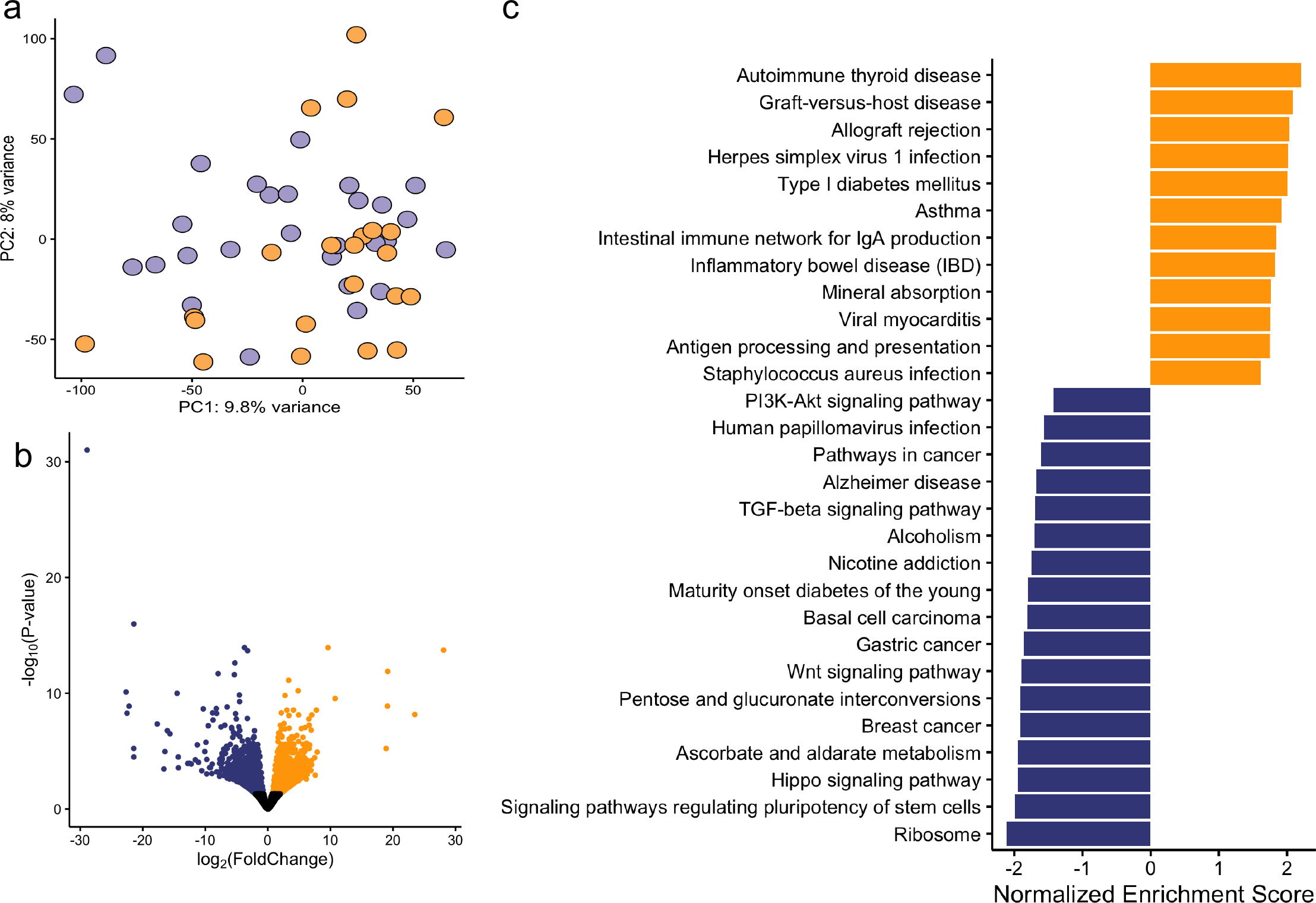
Differentially expressed genes based on BRCA1/2 carrier status. **(A)** Principal components analysis of HGSOC tumors using gene expression data shows a trend towards differences in the tumors based on carrier status, BRCA1/2 carrier (BC) in orange and non-BRCA1/2 carrier (NC) in purple. **(B)** Volcano plot of differentially expressed genes comparing BC tumors vs NC tumors. Significantly up-regulated genes in BC are colored in yellow (padj< 0.05), and significantly down-regulated genes are in blue. **(C)** KEGG gene set enrichment analysis for up-(orange) and down-(blue) differentially expressed genes in BC tumors versus NC tumors.

To connect changes in the methylation of regulatory elements to changes in gene expression, we established a list of 1,782 genes with promoters located within 2 kb of the 135 DMRs that were hypomethylated in tumors from either *BRCA1/2* or *non-BRCA1/2* mutation carriers. We compared this list with the 3,341 DEGs identified in tumors for *BRCA1/2* and non-*BRCA1/2* carriers and identified 68 instances where DMRs showed the predicted inverse correlation between methylation and expression of the nearby gene (Supplementary Table 11) which corresponded to 37 unique regions. These DMRs varied in length from 100bp to 8kb with the number of promoters within 2 kb of a single DMR ranging between one to six. Twenty-five genes were hypermethylated and down-regulated and 41 genes were hypomethylated and up-regulated in tumors from *BRCA1/2* carriers. (Supplementary Table 11). We adapted the software tool ELMER^52,57^ to correlate methylation values in DMRs with the expression of nearby genes; this identified three genes - FGF18, CDK2AP1, and NAGLU - for which methylation and expression were inversely correlated (q<0.05, r= 0.46, 0.44, and 0.48 respectively) (Figure 5c-e). These DMRs correspond to regions that are hypermethylated in tumors from non-*BRCA1/2* carriers, suggesting these regulatory elements suppress the expression of these genes in tumors from *BRCA1/2* carriers.

## DISCUSSION

Whole genome bisulfite sequencing (WGBS) provides a comprehensive genome wide epigenetic landscapes, with coverage of CpGs at several orders of magnitude greater than the array-based approaches largely used up until now to characterize the genome wide methylation status of tumors. In this study, WGBS analysis reported on the methylation status of, on average, 24.6 million CpG sites per tumor. This contrasts with methylation arrays that interrogate highly selected CpG sites, of which the most commonly used have been the Illumina 27K, 450K, and EPIC (850K) arrays that evaluate 0.1%, 1.5% and 3.0% of CpGs in the genome, respectively. The current study is the first to use WGBS to comprehensively map CpG methylation and the transcriptome in matched primary HGSOCs and tumor recurrences arising post chemotherapy in the same patient.

Data are publicly available from genome wide methylation analyses of primary HGSOCs using array based methods. We used data from a 450K methylation array analysis of 20 primary tumors from *BRCA1/2* carriers and 60 primary tumors from *non-BRCA1/2* carriers^21^ to map 3,322 probes that overlap DMRs identified in our WGBS analysis of tumors from *BRCA1/2* and non-*BRCA1/2* carriers. We observed a high concordance (correlation of beta values >0.85) in the methylation level of individual CpGs between WGBS and array measurements of beta values at these sites. Moreover, 749 of the 3,322 probes from 447 unique DMRs were significantly differentially methylated between tumors from *BRCA1/2* and non-*BRCA1/2* cases, with 635 (84.8%) probes showing a concordant direction of effect (**Supplementary Table 12)**. These analyses indicate that WGBS and array based methylation data provide consistently similar readouts for specific probes included on arrays. We were unable to perform a similar correlative analysis using data from the analysis of 613 HGSOCs from TCGA (52 *BRCA1/2* and 561 *non-BRCA1/2* tumors) for which Illumina 27K array data were available due to the low genomic coverage of this array; only one CpG probe (cg21557231) overlapped with DMRs identified in our comparison of *BRCA1/2* and non-*BRCA1/2* tumors. This probe was also differentially methylated in the TCGA data^22^.

TCGA methylation analysis of HGSOCs compared to full thickness fallopian tube tissues described 168 epigenetically silenced genes of which 29 in total, and 4 of the 15 top ranked genes (*AMT, LDHD, CFTR* and *BANK1*) showed significantly reduced expression in non-*BRCA1/2* tumors analyzed in our study. Our data also identified some genes that are differentially methylated in HGSOCs that have been reported by others. In particular *HOXA9* was found to be methylated in up to 95% of ovarian cancers in a study of 80 primary tumors from Montavon and colleagues^58^ and was significantly downregulated in tumors from *BRCA1/2* carriers in our study. Also, *MYO18B* inactivation has been reported in chemoresistant HGSOCs; this gene was significantly downregulated in tumors from *BRCA1/2* carriers from our study^59^.

This is the first study to comprehensively map PMDs genome wide in HGSOCs using WGBS. The data are consistent with studies of the PMD architecture of other cancers and tissues^9,13,14,30,60^. We identified a common set of PMDs, encompassing 15% of the genome that may show specificity to HGSOC. Within these PMDs CpG islands and other functional genomic elements were highly methylated and genes were expressed at lower levels compared to those located outside PMDs^9,13,14^. However, while the distribution of PMDs in primary and recurrent tumors was relatively heterogeneous, there was a strong and highly statistically significant enrichment for genes involved in cancer development, and more specifically genes that are differentially expressed and used to stratify HGSOCs into different molecular subtypes, located outside PMDs. We postulate that the critical nature of these genes in both normal cell function and tumor development requires these genes to be active, both spatially and temporally in their differentiation into specific tumor phenotypes. PMD hypomethylation was a central feature of most of the variation in methylation we observed across our samples. It was only when we masked these regions that we found differences in methylation status between tumors from *BRCA1/2* carriers and *non-BRCA1/2* carriers. The dominance of the PMD signal is an important factor when evaluating tumor methylomes. However, caution needs to be applied to these analyses since masking out PMD regions may result in important focal DMRs remaining undetected.

Previous studies have compared the molecular features of primary ovarian cancers (including methylation) with metastatic tumors or cells from ascitic fluid to look for molecular biomarkers that may represent novel therapeutic targets for chemoresistant disease^21,22,61–63^. Generally, these studies suggest there is an accumulation of somatic changes as tumors metastasize or in recurrences after chemotherapy^21,62^. However, specific mutations associated with tumor metastasis have not been observed across patients in the modestly sized cohorts studied to date. For example, Patch et al^21^ described *ABCB1* promoter fusion in the ascites of 2 out of 15 patients with disease relapse. Whole exome sequencing of 23 pre- and post-chemotherapy exposed primary tumors identified a small number of somatic mutations (between 0-90) that are unique to post-chemotherapy tumors, but none that were observed in more than one patient^62^. Fang et al.^61^ used 450K methylation array analysis to establish a methylation signature of CpGs at promoters for 94 genes in tumor and ascites samples from patients after treatment with cisplatin and the hypomethylating agent guadecitabine that predicts resensitization to platinum^61^. The use of ascites samples is limited because cell populations from ascites are highly heterogeneous and it is not known what proportion of these cells have the ability to seed and grow as solid metastases. While ours and other studies have profiled enriched neoplastic components of solid tumors, the contribution of stromal and other cell contaminants to bulk genomic profiling may have introduced additional heterogeneity. Differences in study design, tissue type analyzed and methylation platforms used across different studies likely contribute to the lack of replication in the data between studies.

Across patients, we unexpectedly found few features in the methylome and transcriptome that were specific to tumor recurrence. Instead, methylation and gene expression signatures were consistently preserved in recurrences relative to the primary tumor from the same individual, with little evidence of an accumulation of additional and novel methylation and transcriptomic changes in the recurrent tumors. This perhaps indicates that the genomic changes required to promote chemo-resistance are established early in primary tumor growth and persist to dominate the clonal population in the primary and subsequent recurrent tumors. This is consistent with a previous study of somatic mutations identified in multiple primary and metastatic samples from seven ovarian cancers, which found complex patterns of both monoclonal and polyclonal seeding of metastatic sites and predicted a lack of selective pressures after treatment with combination chemotherapy^64^. Taken together these studies suggest that many patients have clones of chemoresistant disease at the time of diagnosis; while the bulk of disease responds to platinum-based therapies, a proportion of tumor cells persist through chemotherapy to seed recurrent tumor growth within the peritoneal cavity.

Our WGBS analysis indicates that there are differences in the methylation profiles between patients with and without germline *BRCA1/2* mutations, indicating that hypomethylation is a feature of *non-BRCA1/2* associated tumors. This adds to the growing body of evidence that non-*BRCA1/2* associated ovarian cancers develop along different molecular pathways to ovarian cancers from *BRCA1/2* carriers. Recent studies have suggested that foldback inversions may be drivers of HGSOC development in non-*BRCA1/2* carriers resulting in unique mutational processes that do not correlate with any of the different molecular subtypes described for HGSOC by TCGA^19,22^. Our data support the findings of studies that have identified several notable genes significantly differentially expressed in non-*BRCA1/2* compared to *BRCA1/2* ovarian cancers including: *EIF3CL* which regulates a cluster of metastasis-promoting genes via *STAT3* and acts as a mediator of immune cell evasion^65^ and *CFTR* overexpression (also reported by TCGA) which is known to increase cell invasion, proliferation and adhesion in ovarian cancers^66^, and is highly expressed in the fallopian tube secretory epithelial cells from which many HGSOCs arise^67^.

In conclusion, we have described the first comprehensive analysis of methylation landscapes generated by WGBS in HGSOCs and their recurrences after chemotherapy, and the first comparison in patients with and without germline *BRCA1* and *BRCA2* mutations. This study highlights the molecular heterogeneity that exists amongst HGSOCs and provides the first evidence that this heterogeneity extends to chemoresistant, recurrent disease. We have demonstrated there is an absence of common methylation signatures or specific methylation biomarkers that would indicate common mechanisms and underlying biology shared across patients associated with disease recurrence or chemoresistance. This observation was replicated using whole transcriptome profiling in the same primary-recurrent tissue specimens. Taken together, these data suggest that the methylation and transcriptomic changes required to survive first line chemotherapy and seed recurrent tumors may already be present in the primary tumor, rather than induced as a result of exposure to chemotherapy. Given these findings the continued testing of highly toxic demethylating agents to treat recurrent cancers may not be justified, and alternate drug targets that are effective in treating these tumors are needed. The most significant methylation and/or transcriptomic variations were observed when we compared primary tumors with and without *BRCA1* or *BRCA2* mutations. The improved survival and disease free interval in patients with *BRCA1* or *BRCA2* mutations has been attributed to their improved response to platinum based chemotherapy, and we have identified extensive differences in the methylome and transcriptome between these groups that likely contribute to these differences.

---

**Supplementary Figure 1.**
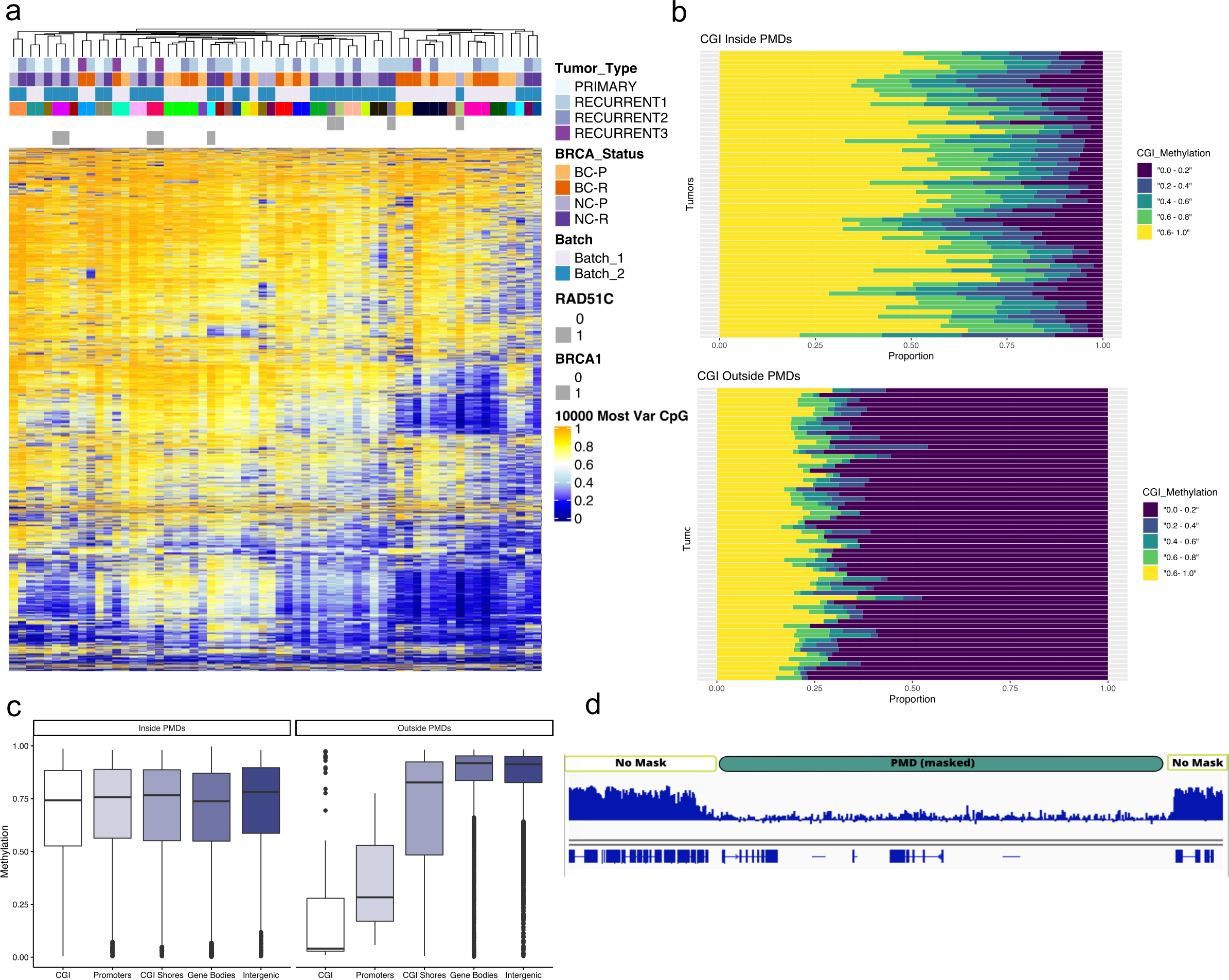
HGSOC tumors show a high degree of heterogeneity. **(A)** Heatmap of the most 10,000 variable CpG sites in the genome, clustered by sample (column) shows that tumors do not cluster by germline mutation, or tumor event status, but by patient. **(B)** CGIs within PMDs are highly methylated, while those outside of PMDs are less methylated. **(C)** Functional elements in the genome are highly methylated when they fall within PMDs. **(D)** Illustration of PMD-masking strategy prior to calling DMRs. PMDs were identified as described in methods, and then those genomic regions were masked out from analysis.

**Supplementary Figure 2.**
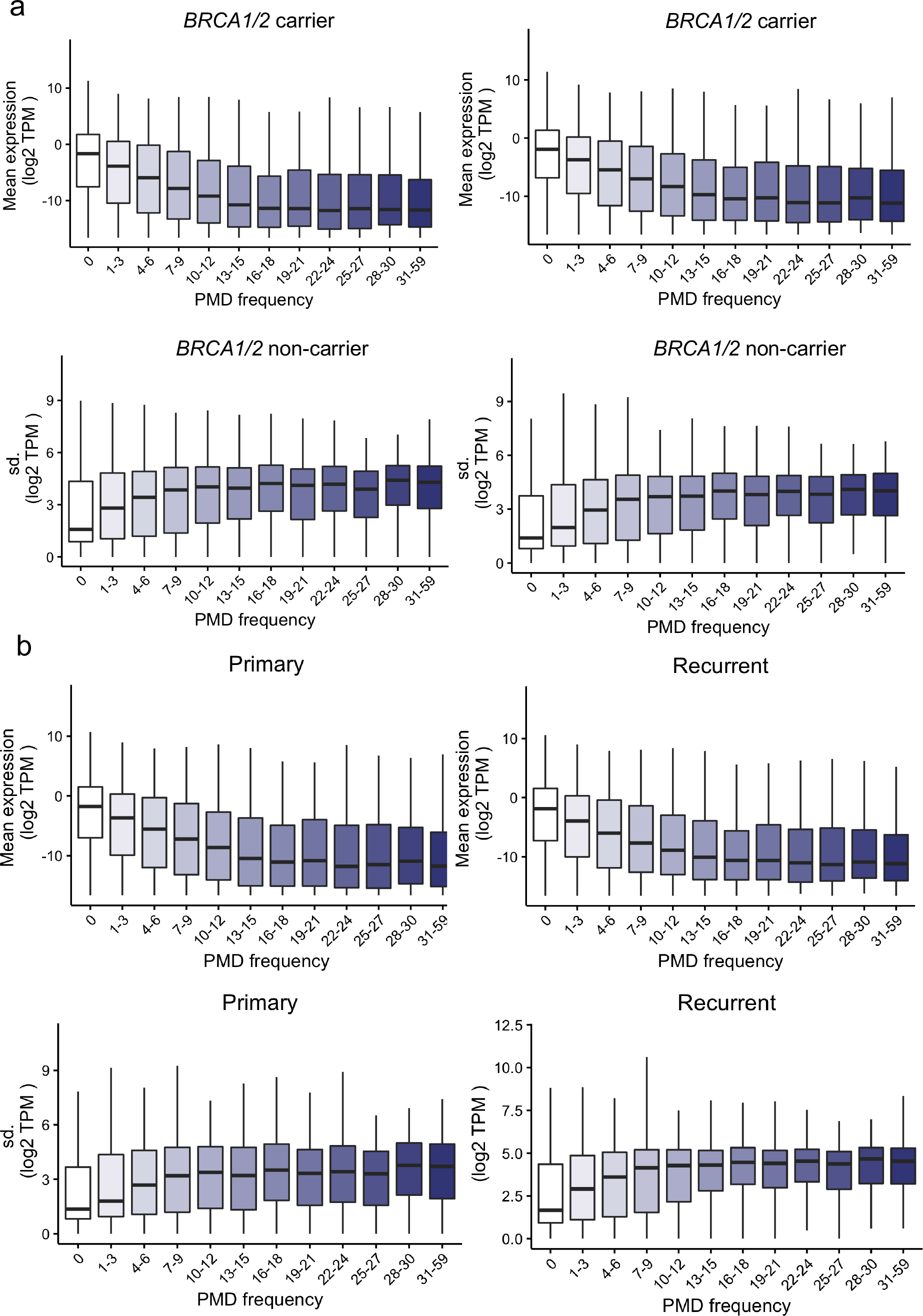
The expression and variability of genes within PMDs in HGSOC tumors. **(A)** Genes frequently within PMDs are expressed at a lower level but are more variable in their expression level than those rarely in PMDs, and this is observed in tumors from BC and NC **(A)**, and equally in primary and recurrent **(B)** tumors.

**Supplementary Figure 3.**
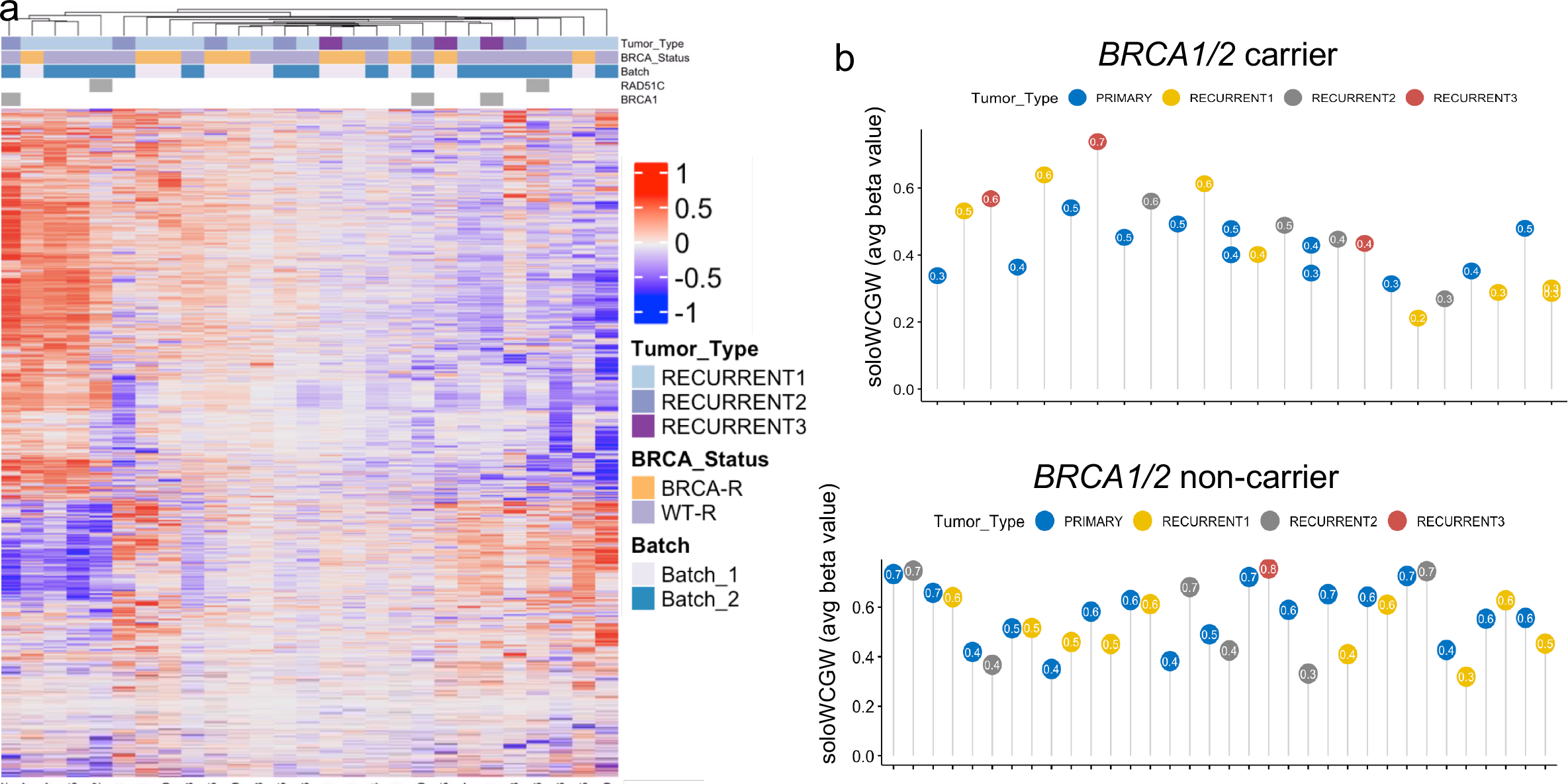
Primary to Recurrent Tumor Progression. **(A)** Heatmap of DMRs from Primary vs Recurrent analysis, (plotted as the delta or change in methylation level between the primary and recurrent tumor) showed variable methylation in the same regions across our tumor sets. Other DMRs indicated relatively no change between primary and recurrent tumors (white regions on heatmap), indicating stability of methylation profile after chemotherapy. **(B)** Average of soloWCGW found within ovcaPMD. Two points on the same stem indicates tumor samples were acquired at the same time.

**Supplementary Figure 4.**
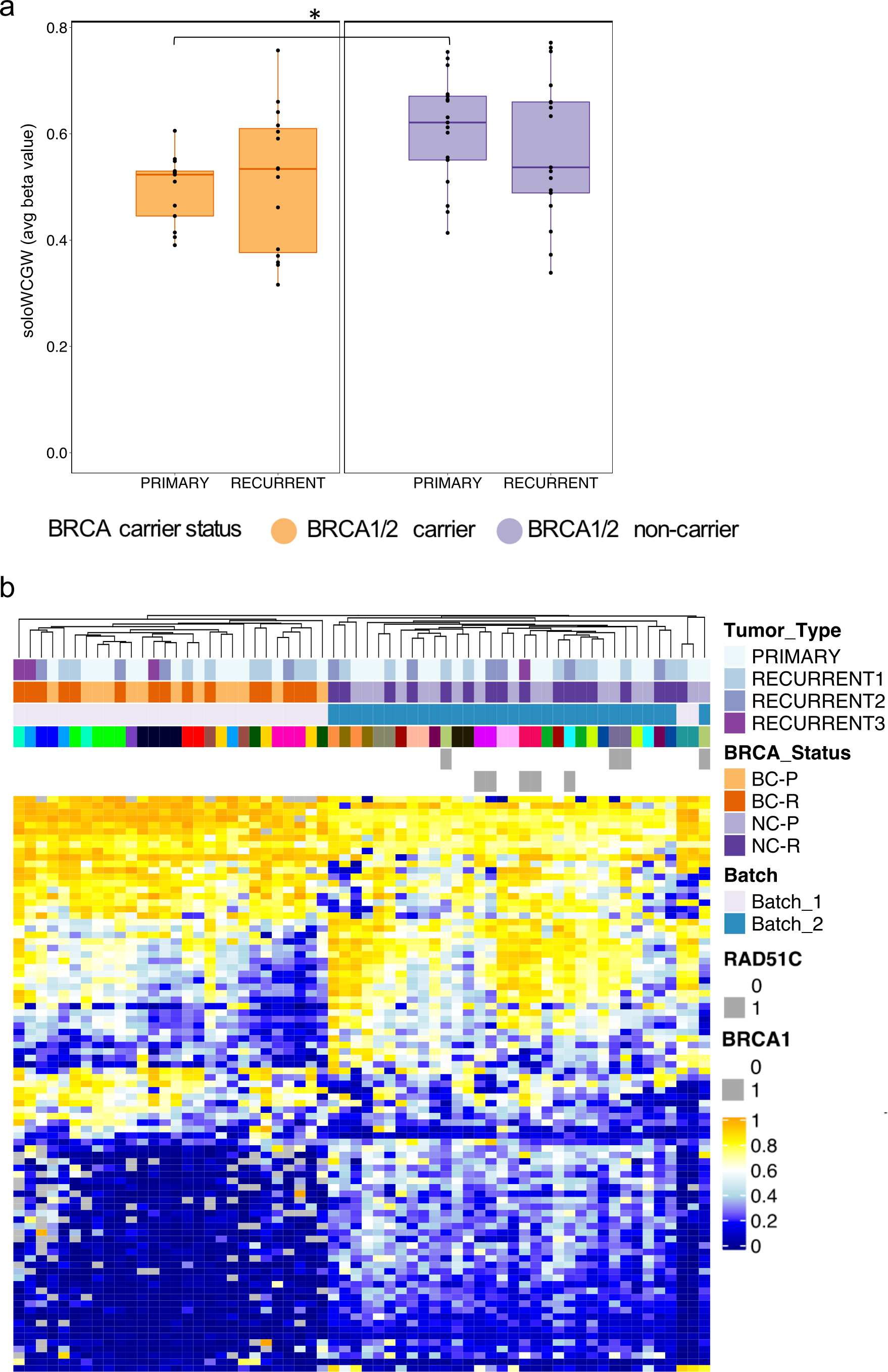
*BRCA1/2* carrier vs *BRCA1/2* non-carrier comparisons. **(A)** *BRCA1/2* non-carrier tumors have significantly higher methylation at soloWCGW sites within PMDs. **(B)** Clustering of tumors based on methylation level at 135 DMRs shows methylation levels were preserved within patients.

**Supplementary Figure 5.**
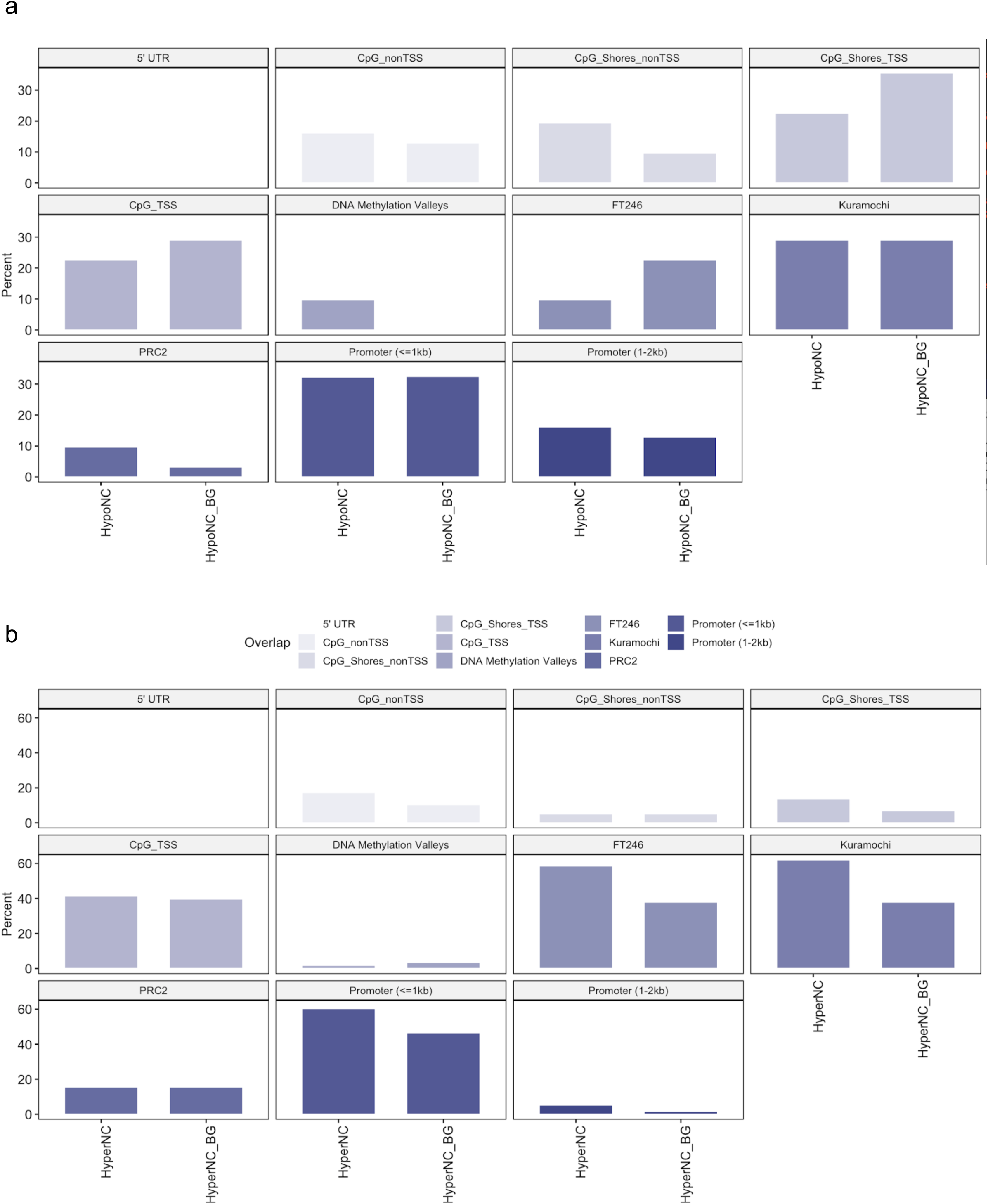
Enrichment of DMRs identified in *BRCA1/2* carrier vs *BRCA1/2* non-carrier tumors. **a)** DMRs hypomethylated in *BRCA1/2* non-carrier tumors. **b)** DMRs hypermethylated in *BRCA1/2* non-carrier tumors.

**Supplementary Figure 6.**
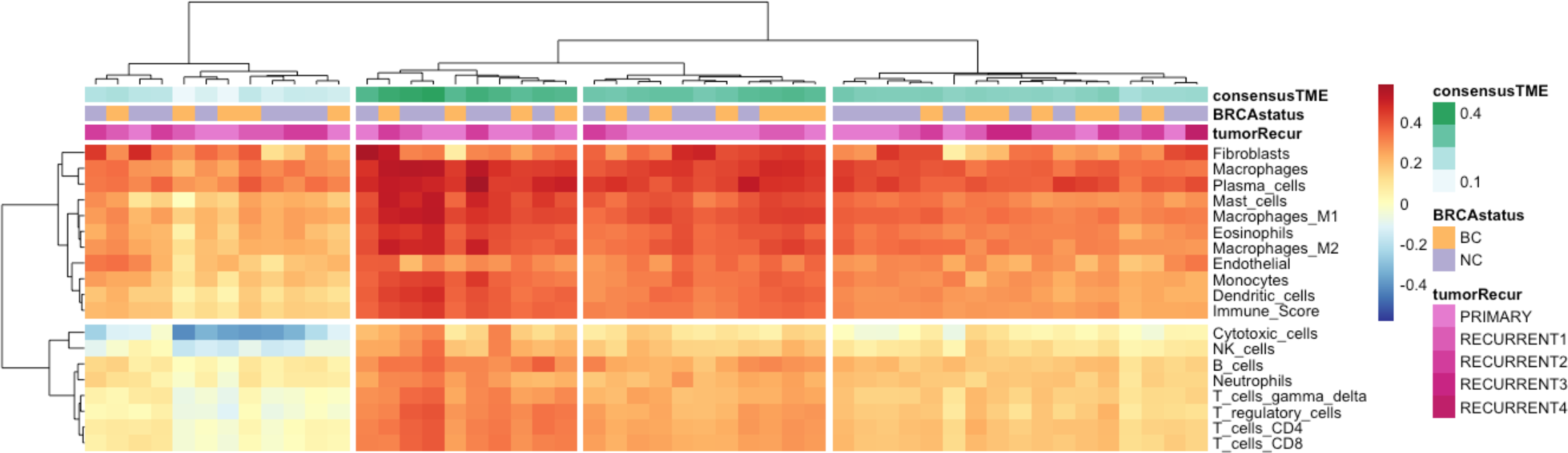
Purity estimates using RNA-Seq data. ConsensesTME cell type estimations identified four clusters of samples defined by a gradient of T cell, B cell Natural Killer cell composition. ConsensusTME clusters did not correlate with *BRCA1/2* status, primary or recurrent status, RNA-Seq library quality, or clinical parameters.

